# A novel SOD1-dependent mechanism for the iron-induced production of toxic SOD1 and oxidative stress that initiates ALS

**DOI:** 10.1101/018846

**Authors:** Liangzhong Lim, Jianxing Song

## Abstract

Free iron is highly toxic and the blood-derived iron initiates early motor-neuron degeneration upon breakdown of blood-spinal cord barrier. Iron is currently known to trigger oxidative stress by Fenton chemistry but no report implies that iron manifests its toxicity through CuZn-superoxide dismutase (SOD1), the central antioxidant enzyme in all human tissues that carries >180 ALS-causing mutations. Here, by NMR we show that Zn^2+^ play an irreplaceable role in the maturation of the nascent hSOD1, and further decipher for the first time that out of 11 other cations only Fe^2+^ has the Zn^2+^-like capacity to induce folding to form the Fe^2+^-bound hSOD1. This acts to reduce or even block the maturation of wild-type and ALS-causing mutant hSOD1, consequently trapping SOD1 in toxic forms and provoking oxidative stress. Our study establishes a novel SOD1-dependent mechanism for iron to manifest cellular toxicity that contributes to pathogenesis of neurodegenerative diseases, aging and even more.

**One Sentence Summary:** Our study establishes a novel SOD1-dependent mechanism for iron to manifest toxicity contributing to neurodegenerative diseases, aging and even more.

---

Amyotrophic lateral sclerosis (ALS) is the most prominent adult motor-neuron disease, clinically characteristic of progressive motor-neuron loss in the spinal cord, brainstem, and motor cortex, which leads to paralysis and death within a few years of disease onset. ALS was first described in 1869, which affects approximately 1-2 per 100,000 people worldwide (1-3). Most ALS cases are sporadic (90%) whereas 10% are familial ALS. Accumulation of iron within CNS was recently identified to be a biomarker of ALS (4-6). Very recently, it has been revealed that in the SOD1^G93A^ mice, the breakdown of blood–spinal cord barrier (BSCB) in the early ALS disease phase led to accumulation of blood-derived iron in the spinal cord, which initiates ALS by triggering early motor-neuron degeneration through iron-induced oxidant stress (7).

Higher eukaryotic aerobic organisms including humans cannot exist without oxygen, but oxygen is inherently dangerous to their existence because of oxidative stress (8-11). Oxidative stress is caused by reactive oxygen species (ROS) arising as by-products of aerobic metabolism, by which superoxide radical is formed in the greatest amount in the metabolism of molecular oxygen. The cell combats harmful ROS with multi-faceted anti-oxidant mechanisms, including catalytic removal of reactive species by enzymes such as superoxide dismutases (SOD), which removes superoxide radicals by catalyzing a dismutation reaction where one superoxide radical is oxidized to form dioxygen and another is reduced to hydrogen peroxide (12-14). Whilst superoxide and hydrogen peroxide are not highly reactive, they can further react to form highly reactive oxidants, such as hydroxyl radical (^.^OH), which are extremely reactive and can cause various damages to proteins, lipids and DNA, thus leading to cell death (8-14). These reactions can be radically accelerated in the presence of reduced metal ions, such as ferrous (8,9).

Three superoxide dismutase isoenzymes are present within human cells, out of which CuZn-superoxide dismutase (hSOD1) is ubiquitously expressed in all tissues and primarily localized in the cytosol, the intermembrane space of mitochondria and the nucleus. hSOD1 is the most abundant, comprising ∼1% of total protein in neurons (12-14). In 1993, hSOD1 was identified to be the first gene associated with FALS (15), and mutations in the SOD1 gene cause the most prevalent form of FALS, accounting for ∼20% of total FALS cases (14-16). Currently, 181 mutations have been identified within the 153-residue hSOD1 that are linked to ALS (http://alsod.iop.kcl.ac.uk/) but their underlying mechanism for triggering ALS pathogenesis remains elusive (16).

Previous studies have established that the monomeric nascent hSOD1 (Figure 1A) folds into the mature form, which is a homodimeric enzyme of remarkably high stability, with each subunit folding into an eight-stranded, Greek-key β-barrel stabilized by an intramolecular disulfide bridge Cys57-Cys146 (12-14). Each subunit holds one copper and one zinc ions in close proximity. While Zn^2+^ is coordinated by His63, His71, His80, and Asp83, while Cu^+^ is ligated by His46, His48, and His120 (12-14) (Figure 1B). However, for the nascent hSOD1 monomer to reach the mature state, a complex multi-step maturation is required in which the critical first step is to recruit zinc, followed by the incorporation of copper and formation of the disulfide bridge catalyzed by copper chaperone for SOD1 (CCS). In contrast to the mature hSOD1, its early species particularly those lacking of the disulfide bridge have been widely shown to have high tendency of aggregation (12-15,17-21). To minimize aggregation, the mutated hSOD1 including the super-stable C6A/C111S mutant were extensively used such as for determining NMR structures lacking of the disulfide bridge (18). Nevertheless, recently it has been found that hSOD1 of the native sequence was highly disordered before metalation and formation of the disulfide bridge *in vivo* (19,20) as well as *in vitro* (21). Even upon excess supplement of zinc, only a population of the unfolded ensemble (Figure 1A) becomes well-folded (Figure 1B), which results in establishment of equilibrium between two forms (19-21).

**Fig. 1.**
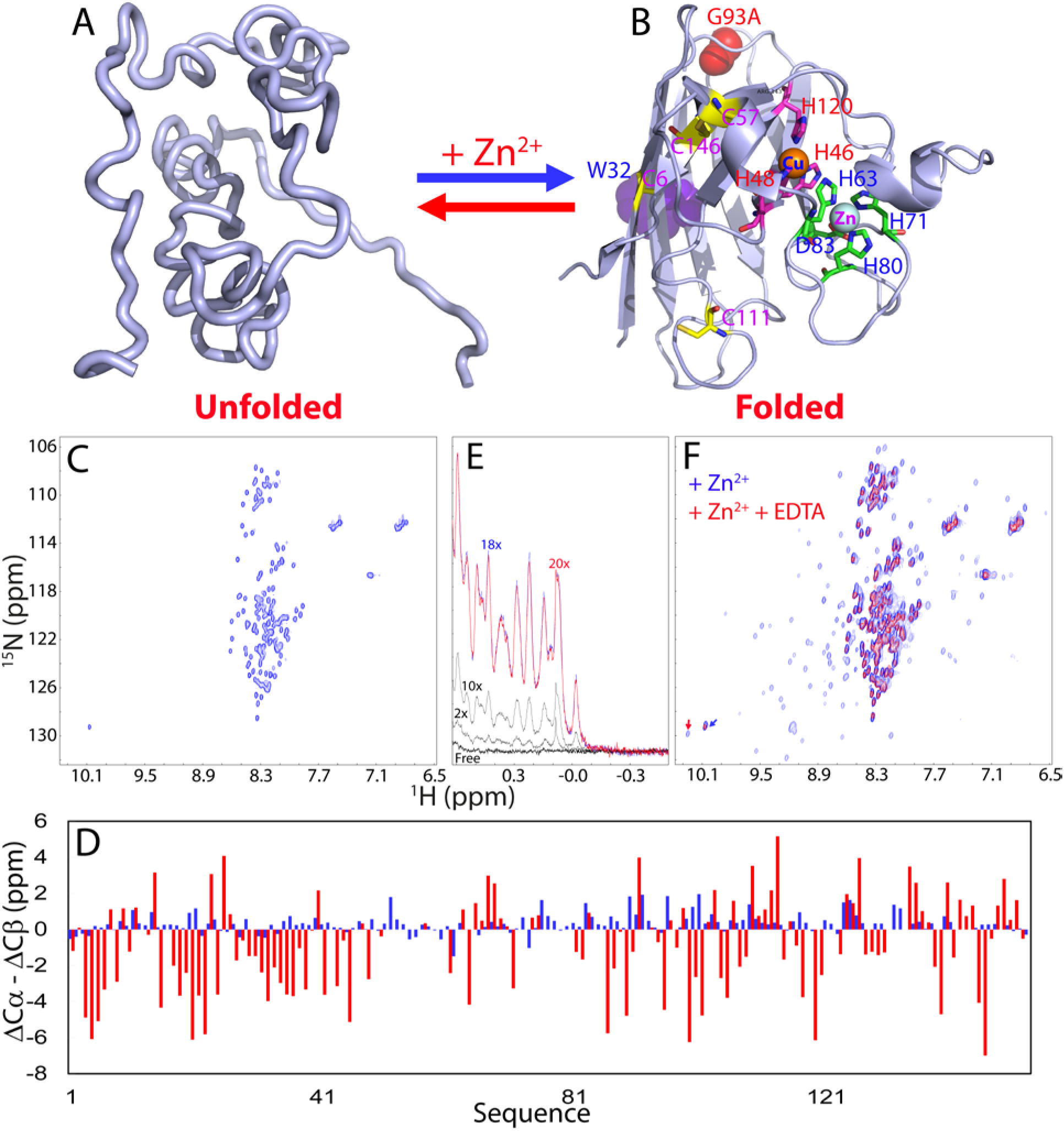
Zn^2+^ plays an irreplaceable role in initiating the hSOD1 maturation. Schematic representation of the unfolded ensemble (A), and mature subunit (B) of hSOD1. The folded hSOD1 is represented by the monomer of the dimeric crystal structure of the human SOD1 (1PU0). Some residues relevant to the present study are labeled. (C) Two-dimensional ^1^H-^15^N NMR HSQC spectrum of hSOD1 of the native sequence without metallation and disulfide bridge. (D) Residue specific (ΔCα-ΔCβ) chemical shifts of hSOD1 without metallation and disulfide bridge (blue); and of the folded form in the presence of zinc (red). (E) Up-field 1D NMR peaks characteristic of the folded SOD1 (-0.5-0.62 ppm) in the presence of zinc at different molar ratios. (F) Superimposition of HSQC spectra of hSOD1 without the disulfide bridge but in the presence of zinc at a ratio of 1:20 (SOD1:zinc) (blue) and that after addition of EDTA at an equal molar ratio of zinc. Blue arrow is used for indicating HSQC peak of the Trp32 ring proton resulting from the unfolded ensemble while red arrow for that from the folded hSOD1.

So far, only zinc is known to own the capacity to trigger the establishment of the conformational equilibrium of hSOD1. In the present study, by high-resolution NMR characterization, we aimed to assess the role of 11 other cations in the crucial first step of the hSOD1 maturation, which include 7 (Na^+^, K^+^, Ca^2+^, Mg^2+^, Mn^2+^, Cu^2+^, and Fe^2+^) essential for humans (22), 3 (Ni^2+^, Cd^2+^ and Co^2+^) previously used to mimic zinc, as well as neurotoxic and environmentally abundant Al^3+^, by evaluating whether they have the zinc-like capacity to trigger the folding of the unfolded hSOD1. Very unexpectedly, while most cations show no such ability, Fe^2+^ was identified for the first time to have the Zn^2+^-like capacity to induce folding to form well-folded Fe^2+^-bound hSOD1. This decodes that Fe^2+^ is able to manifest its cellular toxicity through a SOD1-dependent mechanism.

## RESULTS

### 1. Zinc is essential to initiate the maturation of the nascent hSOD1

Here to achieve high-resolution NMR studies, we have extensively screened and successfully found the optimized solution condition which allow the preparation of high-concentration samples without aggregation (up to 500 μM) of hSOD1 of the native sequence. Consistent with recent *in vivo* (19,20) and *in vitro* results (21), hSOD1 of the native sequence is unfolded without metalation and disulfide bridge, as clearly reflected by its poorly-dispersed HSQC spectrum (Figure 1C), which thus mimics the nascent hSOD1. By analyzing a large set of three-dimensional NMR spectra, we succeeded in achieving its NMR assignments and Figure 1D presents its (ΔCα–ΔCβ) chemical shift, which is an indicator of the residual secondary structures in disordered proteins (21,23). The small (ΔCα–ΔCβ) chemical shifts for most residues clearly indicate that the nascent hSOD1 is an ensemble of highly disordered conformations in solution, which retains no native β-sheet structure. Furthermore, hNOEs, a sensitive indicator of the backbone motions on the ns–ps time scale (21,23), are small for all residues with an average value of –0.10 (Figure S1A). In particular, more than half of the residues have negative hNOEs, clearly indicating that this unfolded ensemble is very flexible with highly unrestricted ns–ps backbone motions. We also collected ^15^N backbone CPMG relaxation dispersion data and found no significant dispersion, indicating that under this condition, the highly-disordered hSOD1 ensemble is lacking of significant exchanges on the μs–ms time scale with significant chemical shift differences (24).

By contrast, once Zn^2+^ was added, a well-folded population was formed as indicated by the manifestation of up-field 1D peaks (Figure 1E), as well as many well-dispersed HSQC peaks (Figure 1F), which are characteristic of the folded population. A stepwise addition of Zn^2+^ triggers gradual increases of the folded population which is largely saturated at a ratio of ∼1:18 (hSOD1:zinc). Interestingly, even with the ratio reaching 1:40, the unfolded and folded conformations still remain coexisting, as clearly reflected by the presence of two HSQC peaks (one characteristic of unfolded and another of unfolded) from the aromatic ring of Trp32 (Figure 1F). This implies that even excess presence of zinc is insufficient to completely shift the unfolded ensemble to the folded form, consistent with recent *in vivo* (19-20) and *in vitro* (17,21) observations. Amazingly, we also observed this phenomenon on the N-terminus of another ALS-causing protein TDP-43, which encodes equilibrium of the folded and unfolded ubiquitin-like domain (25).

By analyzing triple-resonance NMR spectra acquired on a double-labeled sample in the presence of Zn^2+^ at 500 μM, we have achieved sequential assignments of the folded form of the Zn^2+^-bound hSOD1, and its (ΔCα–ΔCβ) chemical shifts were shown in Figure 1D. The very large (ΔCα–ΔCβ) chemical shifts for most residues clearly reveal that Zn^2+^ indeed has the unique capacity in triggering the formation of the well-folded structure, consistent with recent in-cell results (19,20). Previously, a Zn^2+^-bound structure of hSOD1 with super-stable mutations (C6A/C11S) was determined by NMR (18), and Figure S1B presents (ΔCα–ΔCβ) chemical shifts of this super-stable mutant (BMRB Entry 6821) and our folded hSOD1 of the native sequence. The results showed that except for several residues close to the mutation site C6 and C111, the (ΔCα–ΔCβ) chemical shifts of both forms are highly similar for the assigned residues. This provides strong evidence that upon binding to Zn^2+^, both of them adopt highly similar structures despite having difference of two residues.

The well-dispersed peaks disappeared upon adding EDTA at an equal molar concentration of Zn^2+^ (Figure 1F), confirming that the formation of the folded species is a Zn^2+^-specific effect. Furthermore, we generated the H80S/D83S mutant which was previously shown to have severely-abolished capacity in zinc-binding (26). As shown in Figure S2A, this mutant is also highly-disordered like the wild-type hSOD1, as evident by its poorly-dispersed HSQC spectrum with most peaks superimposable to those of the wild-type except for those of mutated residues and several residues close to the mutation sites in sequence. Indeed, upon addition of zinc at a ratio of 1:10, only very minor change was observed for up-field peaks (Figure S2B) and the changes were mostly saturated at 1:30. However, even at a ratio of 1:40, the majority of well-dispersed HSQC peaks characteristic of the well-folded hSOD1 is not detected (Figure S2C). On the other hand, several relatively-dispersed HSQC peaks of the mutant in the presence of Zn^2+^ are mostly superimposable to those of the Zn^2+^-bound wild-type hSOD1 (Figure S2D). The results together indicate that the mutations significantly reduce the zinc-binding capacity as demonstrated by the crystal structure lacking of Zn^2+^ (26). Consequently, for this mutant, even the excess supplement of Zn^2+^ could only trigger the formation of the partially-folded form.

### 2. Most cations show no capacity in triggering the folding

Previously, other metal ions have been demonstrated to be capable of substituting either zinc or/and copper ions in the mature SOD1 (12-14) but it remains unknown whether they have the Zn^2+^-like capacity in triggering the folding of the unfolded hSOD1. In order to address this, we conducted HSQC titrations of the unfolded hSOD1 with 11 cations with a ratio of hSOD1:ion reaching up to 1:40. The results revealed that most cations show no detectable capacity in triggering the folding. For example, Ni^2+^, Cd^2+^, Mn^2+^ and Co^2+^, which were used to substitute Cu^2+^ or Zn^2+^ in the matured SOD1, only induced very minor shifts of several HSQC peaks but have no detectable capacity in triggering the folding (Figure S3). This highlights the irreplaceable role of Zn^2+^ in initiating the hSOD1 maturation (17,19,20). Indeed, previous folding studies have deciphered that zinc modulates the entire folding free energy surface of hSOD1 (27). So an intriguing question arises: why does the wild-type hSOD1 still retain equilibrium between the folded and unfolded forms even in the excess presence of Zn^2+^? Previous studies revealed a surprising fact that on the one hand, to achieve copper-load and disulfide formation catalyzed by hCCS, the hSOD1 needs to populate the folded form bound to Zn^2+^ to some degree. Otherwise, the efficacy of maturation was low, as exemplified by the G93A-hSOD1 mutant (13,17,19,20,28,29). On the other hand, the interaction between hCCS and folded hSOD1 cannot be too stable but has to be dynamic/transit. If the hSOD1-hCCS complex is too stable, as observed on the H46R/H48Q mutant, the efficiency of the hCCS-catalyzed maturation will also be reduced or even abolished (29). So the request for dynamic/transit interactions between hSOD1 and hCCS appears to be elegantly fulfilled by the dynamic nature of the immature hSOD1, which at least partly results from the co-existence of the unfolded and folded population in the presence of zinc, as we recently deciphered for the TDP-43 N-terminus (25).

Remarkably, Cu^2+^ induced much more extensive shifts of HSQC peaks than Zn^2+^ (Figure S4A), implying that Cu^2+^ is able to significantly bind the unfolded hSOD1 ensemble. However, Cu^2+^ was only able to trigger the formation of a partially-folded form because most well-dispersed HSQC peaks characteristic of the Zn^2+^-induced folded form were not detectable in the Cu^2+^-induced form (Figure S4A). This result is similar to what was observed on the H80S/D83S mutant induced by Zn^2+^ (Figure S2). Interestingly the Cu^2+^-induced effect is mostly saturated at ∼1:6 (hSOD1: Cu^2+^) (Figure S4B). We also observed that the hSOD1 samples in the presence of Cu^2+^ are prone to aggregation at high Cu^2+^ or/and protein concentrations. The extensive binding to the unfolded ensemble, inability to induce the well-folded form, and induced aggregation by Cu^2+^ may partly account for its high *in vivo* toxicity and also rationalize why the load of copper has to be specifically catalyzed by hCCS to the pre-folded hSOD1 population maintained by binding with Zn^2+^ (17,19,20).

### 3. Fe^2+^ has the capacity in triggering the folding to form the Fe^2+^-bound hSOD1

The most unexpected finding in this study is that except for Zn^2+^, Fe^2+^ is the only one out the of 11 cations that owns the capacity in triggering the folding to form the well-folded Fe^2+^-bound hSOD1. As seen in Figure 2A, addition of Fe^2+^ triggers the manifestation of up-field NMR peaks characteristic of the folded form and the increase in the folded population is mostly saturated at a molar ratio of 1:20 (hSOD1: Fe^2+^). However, a detailed comparison of the intensity and chemical shift of up-field peaks induced by Fe^2+^ to that by Zn^2+^ is impossible as Fe^2+^ has strong paramagnetic effects including effects of relaxation enhancement and pseudo-contact shift (30). Figure 2B presents the superimposition of the HSQC spectra of the Fe^2+^- and Zn^2+^-bound hSOD1 forms respectively. Based on the assignments of the Zn^2+^-bound form, many peaks from the Fe^2+^-bound hSOD1 are highly superimposable to those of the Zn^2+^-bound; while some have significant shifts. Interestingly, as shown in Figure 2C, the peaks with significant shifts are resulting from the residues surrounding the Cu/Zn binding pocket, while the peaks without significant shifts are from those located relatively far away from the Cu/Zn binding pocket. This strongly suggests that the Fe^2+^- and Zn^2+^-bound hSOD1 structures are very similar particularly over the beta-barrel region, and further implies that Fe^2+^ binding site is located close to, or even has overlaps with the Cu/Zn binding pocket. Consequently the residues close to Fe^2+^ showed large shifts for their HSQC peaks due to the pseudo-contact shift effect (30), or/and slightly-different conformations in the Fe^2+^-bound form. Moreover, the up-filed and well-dispersed HSQC peaks also disappeared upon adding EDTA at an equal molar concentration of Fe^2+^, confirming that the formation of the folded hSOD1 is also Fe^2+^-specific effect.

**Fig. 2.**
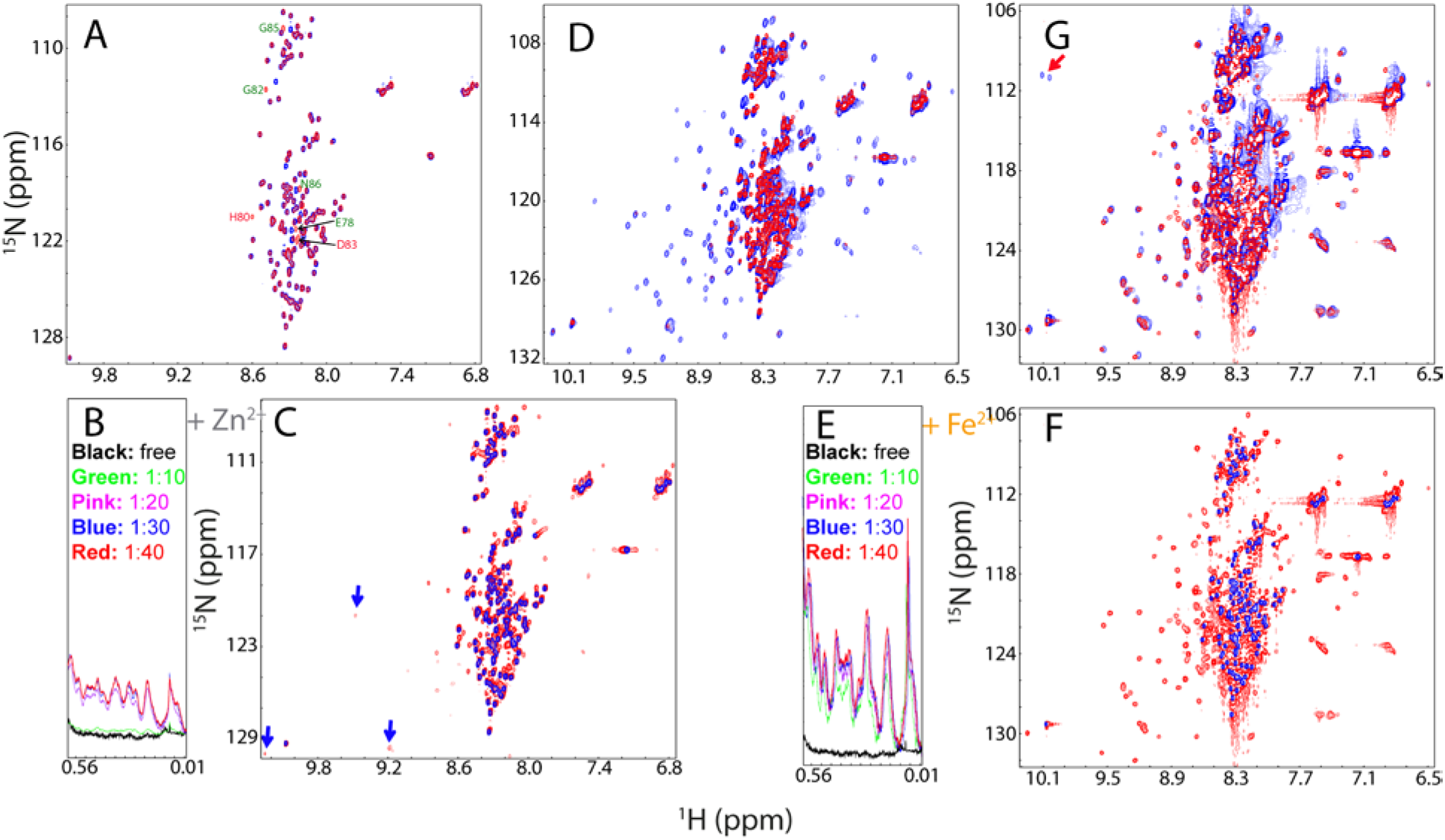
Fe^2+^ has the Zn^2+^-like capacity in triggering the folding to form the Fe^2+^-bound hSOD1. (A) Up-field 1D NMR peaks characteristic of the folded hSOD1 (-0.5-0.62 ppm) in the presence of Fe^2+^ at different molar ratios. (B) Superimposition of HSQC spectra of hSOD1 without the disulfide bridge but in the presence of Zn^2+^ (blue), and Fe^2+^ (red). The labels of the sequential assignments for some well-resolved HSQC peaks are in green if the peaks are largely superimposable but in pink if the peaks are largely shifted in both forms. (C) The monomer structure of hSOD1 (1PU0) with the residues displayed in pink spheres if their HSQC peaks largely shifted in the Zn^2+^- and Fe^2+^-forms, and colored in green if their HSQC peaks almost superimposable. (D) Up-field 1D NMR peaks (-0.5-0.62 ppm) of hSOD1 in the presence of 20x Zn^2+^ (green), 20x Fe^2+^ (blue), addition of 10x Zn^2+^ into the sample in the pre-existence of 20x Fe^2+^ (black); and addition of 20x Zn^2+^ into the sample in the pre-existence of 20x Fe^2+^ (red). (E) Superimposition of HSQC spectra of hSOD1 in the presence of 20x Zn^2+^ (blue), in the presence of 20x Fe^2+^ (green), and in the presence of both 20x Fe^2+^ and 10x Zn^2+^. (F) Superimposition of HSQC spectra of hSOD1 in the presence of 20x Zn^2+^ (blue), in the presence of 20x Fe^2+^ (green), and in the presence of both 20x Fe^2+^ and 20x Zn^2+^. Red arrows are used for indicating Fe^2+^-specific HSQC peaks while blue arrows for Zn^2+^-specific peaks.

We also conducted a competitive experiment between Zn^2+^ and Fe^2+^ by monitoring the changes of both up-field (Figure 2D) and HSQC (Figures 2E and 2F) peaks upon stepwise addition of Zn^2+^ into a hSOD1 sample in the pre-existence of 20x Fe^2+^. As seen in Figure 2D, with addition of 10x zinc, the zinc-specific peak in the 1D spectrum manifested. On the other hand, however, as shown in HSQC spectra (Figure 2E), in the presence of 10x zinc, many HSQC peaks still remain highly similar to those of the Fe^2+^-bound. Upon addition of 20x Zn^2+^, although many peaks become superimposable to those of Zn^2+^-bound form, there are still some peaks more superimposable to those of the Fe^2+^-bound form (Figure 2F). This implies that Fe^2+^ and Zn^2+^ binding-sites are only partly overlapped. Unfortunately, further addition of Zn^2+^ triggers severe broadening of the NMR peaks and aggregation of the sample, thus blocking our further characterization. Nevertheless, the results imply that even for the wild-type hSOD1, the presence of Fe^2+^ at high concentrations interferes in the Zn^2+^-induced formation of the well-folded hSOD1, thus reducing the efficiency of the hSOD1 maturation.

We titrated the H80S/D83S mutant with Fe^2+^ by monitoring the changes of upfield 1D (Figure S2E) and HSQC (Figure S2F) spectra. Although for this mutant, zinc failed to trigger the formation of the well-folded structure (Figure 2C), Fe^2+^ is still able to as evident from the manifestation of up-field peaks mostly saturated at a ratio of 1:20 (Figure S2E), as well as well-dispersed HSQC peaks (Figure S2F), which are mostly superimposable to those of the wild-type (Figure S2G). These results suggest that the Fe^2+^-bound structures are highly similar for both H80S/D83S mutant and wild-type hSOD1.

To explore the relationship of the Fe^2+^-binding site to the Cu^2+^ one, we generated another mutant carrying two ALS-causing mutations, H46R/H48Q whose Cu^2+^-binding capacity is found to be mostly eliminated (29). Very surprisingly, even before metalation and formation of the disulfide bridge, this mutant already has a well-folded population (Figure S5A). This result thus rationalizes the previous observation that even this mutant is completely demetallated, it formed a much more stable complex with hCCS than the wild-type. Surprisingly, the formation of the stable complex acted to block, rather than to enhance further formation of the intra-subunit disulfide bridge catalyzed by hCCS (29). As such, this mutant was found to be trapped in the soluble form lacking of the disulfide bridge, which appeared to be highly toxic for initiating ALS pathogenesis (29).

For this mutant, addition of Zn^2+^ triggers the increase of the folded population which is also saturated at a ratio of 1:20 (Figure S5B). However, it seems that at 1:20, the intensity of up-field peaks (Figure S5B) is lower than that for the wild-type (Figure 1E), consistent with the previous result that this mutant also has reduced zinc-binding affinity (29). Consistent with the 1D result, the intensity of the well-dispersed HSQC peaks increases at a ratio of 1:20 (Figure S5C). Furthermore its HSQC peaks are mostly superimposable to those of the wild-type triggered by Zn^2+^ (Figure S5D), implying that the overall structures are very similar for the mutant and wild-type. Most importantly, Fe^2+^ is again able to trigger the significant increase of the folded population which is also saturated at a ratio of 1:20 (Figure S5E). Similarly, in the presence of Fe^2+^ at 1:20, the intensity of the well-dispersed HSQC peaks significantly increases (Figure S5F), and their HSQC peaks are mostly superimposable to those of the wild-type triggered by Fe^2+^ (Figure S5G). The results together suggest that the Fe^2+^-binding site is also not completely overlapped with the Cu^2+^ one.

We have also assessed the interactions of hSOD1 with Fe^2+^, Zn^2+^ and Cu^2+^ by isothermal titration calorimetry (ITC) measurements. Similar patterns of heat releases are observed for the titrations of the unfolded hSOD1 with three cations (Figure S6). However, as the heat changes are expected to be associated with at least two processes: namely binding of hSOD1 with cations and binding-induced conformational changes, data fitting is not possible to obtain the thermodynamic parameters for only binding.

### 4. Fe^2+^ radically blocks the Zn^2+^-induced folding of G93A-hSOD1

So far, >181 ALS-causing mutations have been identified on the 153-residue hSOD1 but it still remains highly elusive for the mechanism by which these mutations cause ALS. To explore this, we generated the ALS-causing G93A-hSOD1 protein (29) which is also highly disordered like the wild-type as judged by its poorly-dispersed HSQC spectrum (Figure S7). Interestingly, however, in addition to the residues such as Asp92, Val94 and Ala95 which are next to the mutation residue Gly93 in sequence, many residues such as Gln22, Glu24, Asp101 and Asp103 far away from Gly93, also have significant shifts for their HSQC peaks. This is very different from what observed on the H80S/D83S mutant (Figure S2A), implying that even in the unfolded ensemble, the ALS-causing mutation of G93A is able to trigger long-range perturbations on other residues.

Addition of Zn^2+^ is still able to trigger the formation of the folded form of G93A-hSOD1, as evident from the manifestation of up-field 1D (Figure 3A) and well-dispersed HSQC peaks. However, as compared to the titration results of the wild-type SOD1 by Zn^2+^, two significant differences were observed (Figure 3A). First, at the same ratio, the intensity of up-field peaks of G93A is much lower than that of the wild-type hSOD1. Second, the Zn^2+^-induced formation of the G93A folded form will not be saturated even up to 1:60. However, attempts to further add Zn^2+^ triggered severe aggregation of G93A-hSOD1, thus preventing from further characterization at higher ratios of Zn^2+^. Nevertheless, both differences imply that the G93A hSOD1 has a considerably reduced capacity in forming the folded form induced by Zn^2+^.

**Fig. 3.**
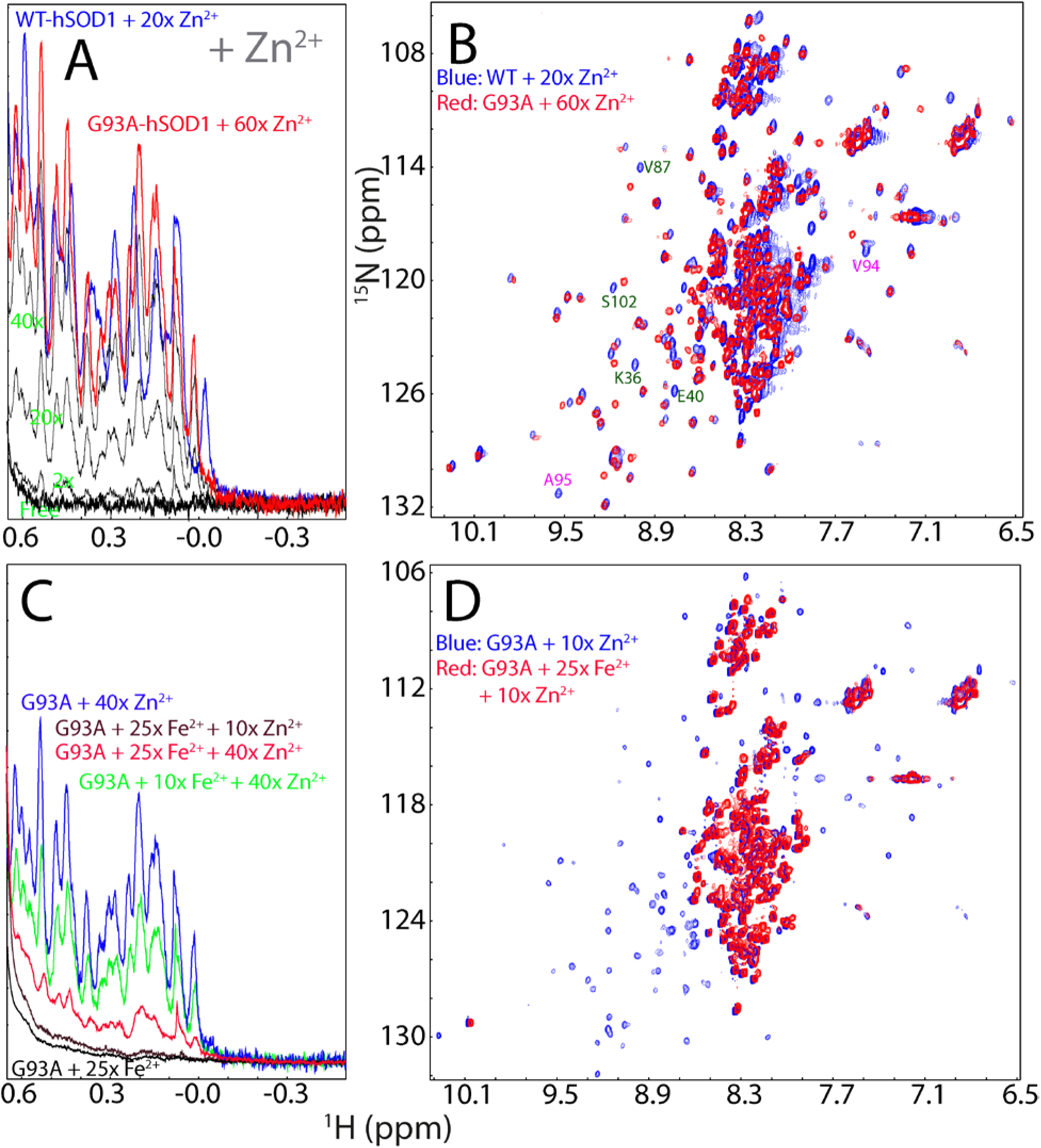
Fe^2+^ significantly reduces the efficiency of the first step of the G93A-hSOD1 maturation. (A) Up-field 1D NMR peaks characteristic of the folded wild-type or G93A hSOD1 (-0.5-0.62 ppm) in the presence of Zn^2+^ at different molar ratios. (B) Superimposition of HSQC spectra of the wild-type hSOD1 in the presence of Zn^2+^ at a molar ration of 1:20 (blue), and G93A hSOD1 in the presence of Zn^2+^ at a molar ration of 1:60 (red). Some residues with significant shifts of their HSQC peaks are labeled (C) Up-field 1D NMR peaks characteristic of the folded G93A hSOD1 (-0.5-0.62 ppm) with a pre-existence of 10x or 25x Fe^2+^, followed by addition of Zn^2+^ at different molar ratios. (D) Superimposition of HSQC spectra of G93A-hSOD1 in the presence of 10x Zn^2+^ without (blue) and with the pre-existence of 25x Fe^2+^ (red).

Amazingly, although the G93A mutation is not located within the metal binding pocket, Fe^2+^ failed to trigger the formation of the folded form. Even in the presence of 80x Fe^2+^, no peaks characterized by the folded form manifested over up-field region (Figure S8A), and in its HSQC spectrum (Figure S8B). On the other hand, the presence of Fe^2+^ led to significant shifts of some HSQC peaks, most of which are from the residues close to the Cu/Zn binding pocket except for Ala4, Leu6, Ile113 and the mutation residue Ala93 (Figure S8C), indicating that Fe^2+^ is able to bind to these residues in the unfolded ensemble of the G93A hSOD1.

hSOD1 accounts for ∼1% of the total proteins in neurons. If assuming that the cellular protein concentration is ∼200 mg/ml, the hSOD1 concentration is 2 mg/ml, or 125 μM. On the other hand, it has been found that in the SOD1^G93A^ mice, the breakdown of blood–spinal cord barrier (BSCB) could result in the availability of free Fe^2+^ in neurons up to 800 ng/mg protein, which is ∼2.86 mM (7). That means that free Fe^2+^ concentration is ∼23 time higher than the hSOD1 concentration in neurons. Therefore, here to mimic the *in vivo* situation, we evaluated the efficiency of Zn^2+^ in triggering the folding of G93A in the pre-existence of Fe^2+^ at ratios (G93A: Fe^2+^) of 1:10 and 1:25 respectively (Figure 3C). Interestingly, even in the presence of only 10x Fe^2+^, the intensity of up-field peaks induced by adding 40x zinc is much lower than that without Fe^2+^ (Figure 3C). The intensity of the HSQC peaks are also much weaker in the presence of 10x Fe^2+^, although most HSQC peaks are still superimposable to those in the presence of 40x zinc without Fe^2+^ (Figure S9). Most strikingly, in the pre-existence of 25x Fe^2+^, addition of 10x zinc is no longer able to induce up-field 1D (Figure 3C), as well as well-dispersed HSQC peaks (Figure 3D). Only upon addition of 40x zinc, the up-field peaks manifested to some degree, but however, those peaks are very weak and broad (Figure 3C). Furthermore, visible aggregates were observed shortly in NMR tube and consequently, no high-quality HSQC spectrum could be acquired on this sample. Although the exact Zn^2+^ concentration in neurons remains to be defined, it was estimated that the Zn^2+^ concentration is only in the 10–30 μM range even upon being released (31), which is not even reaching a ratio of 1 to 1 with hSOD1. As such, upon the breakdown of the blood–spinal cord barrier, because of the binding of Fe^2+^ to the unfolded G93A ensemble, the maturation of the G93A-SOD1 is expected to be significantly retarded in the neurons, consistent with the *in vivo* observations (13,14,28,29).

### Discussion

Oxidative stress, resulting from an imbalance between the production of free radicals and the ability of the cell to remove them, has been increasingly identified to cause various human diseases, particularly neurodegenerative diseases (7-11). In this context, hSOD1 represents a central antioxidant while iron acts as a notorious accelerator. Iron is a double-edged sword: it is the most abundant “trace element” absolutely required for humans’ survival, but high levels of iron quickly lead to cell death. Under the normal conditions, the cellular concentration of free iron is very low (22,32), but under the pathological conditions such as the breakdown of the blood-central nerve system characteristic of neurodegenerative diseases and aging (33,34), the concentration of the blood-derived iron can reach very high (4-7). Indeed, iron is accumulated in various neurodegenerative as well as other diseases but the underlying mechanisms for its toxicity still remain to be fully elucidated (35-40). Currently, iron is known to trigger oxidative stress mainly through its reactivity with peroxide, thus generating the highly reactive hydroxyl radical by Fenton chemistry (8,35-40). So far, there is no report implying that iron might manifest its cellular toxicity by specifically targeting SOD1.

Here, by high-resolution NMR studies, we first demonstrated that the hSOD1 of the native sequence is highly disordered, and Zn^2+^ plays an irreplaceable role in initiating its maturation by triggering the formation of the folded population bound to Zn^2+^. Previous studies revealed that both overall folded structure and local conformations/dynamics of hSOD1 maintained by Zn^2+^ are essential for further load of copper and oxidation to form the disulfide bridge Cys57-Cys146 as further catalyzed by hCCS (17,19,20). Most strikingly, in this study, for the first time, we decode that out of 11 other cations, only Fe^2+^ has the Zn^2+^-like capacity to induce the folding to form the Fe^2+^-bound hSOD1, which is most likely unable to complete further load of copper and formation of the disulfide bridge, probably partly due to the differences of the Fe^2+^- and Zn^2+^-bound hSOD forms in local conformations/dynamics, or/and to the fact that Fe^2+^ has the binding site partly overlapped with that of copper. Indeed, so far, there has been no report on detecting the Fe^2+^-bound active hSOD1 *in vivo*. Therefore, the presence of Fe^2+^ at high concentrations reduces the efficiency of the Zn^2+^-induced maturation of both wild-type and ALS-causing mutant hSOD1. Recently the failure or even reduced efficiency of the maturation has been proposed as a common mechanism for trapping the ALS-causing mutants in the highly-toxic species without the disulfide bridge, which are also prone to aggregation (13,29,30). Consequently, our finding establishes a novel SOD1-dependent mechanism for the iron-induced production of oxidative stress, as well as highly toxic forms of the mutant and wild-type hSOD1.

In the context of numerous previous results, we propose here a mechanism by which Fe^2+^ targets hSOD1 to provoke oxidative stress as well as to create toxic hSOD1 forms (Figure 4). More specifically, the hSOD1 of the native sequence before metalation and disulfide bridge formation exists as a disordered ensemble, thus mimicking the nascent hSOD1 (Figure 4A). Upon supplement of Zn^2+^, a conformational equilibrium is established in which the folded population is formed (Figure 4B), thus ready for further interacting with hCCS (Figure 4C) to reach the mature and active hSOD1 with copper loaded and the disulfide bridge formed capable of catalyzing the dismutation reaction (Figure 4E). However, if in the presence of Fe^2+^ at high concentrations, a population of hSOD1 becomes the Fe^2+^-bound form, which has an overall architecture similar to the Zn^2+^-bound (Figures 4F and 4G), but unsuitable for further load of copper and formation of the disulfide bridge catalyzed by hCCS, As a consequence, the Fe^2+^- bound hSOD1 form without stabilization by the disulfide bridge acquires high toxicity as found with other ALS-causing mutants including G93A (28) and H46R/H48Q (29), by such as interacting with membranes of organelles (21,41) or/and become aggregated to form the iron-containing hSOD1 inclusion (Figure 4H) extensively observed in ALS patients.

**Fig. 4.**
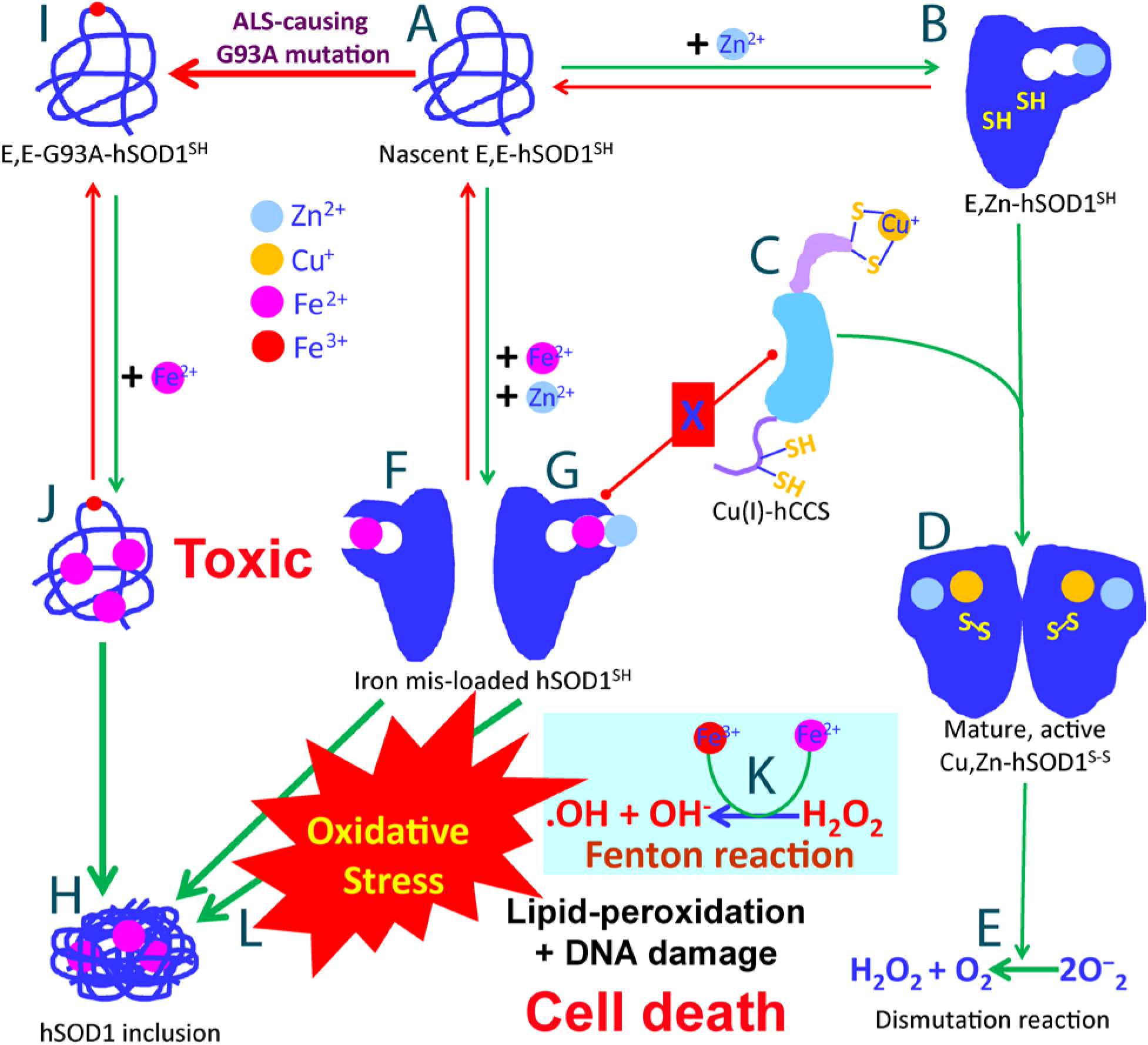
A SOD1-dependent mechanism by which Fe^2+^ provokes oxidative stress and traps the wild-type and ALS-causing mutant hSOD1 in the toxic forms.

For G93A-hSOD1 (Figure 4I), even without the excess presence of Fe^2+^, the efficiency of the Zn^2+^-induced maturation has been demonstrated to be significantly reduced (13,28), likely owing to the reduced folding capacity induced by Zn^2+^ as we reveal here (Figure 3A). This reduction is expected to provoke oxidative stress which may subsequently trigger the breakdown of blood–central nerve system barrier as previously demonstrated on the SOD1^G93A^ mice (42,43). Upon the breakdown, the concentration of the blood-derived Fe^2+^ could become as high as ∼25 times of the hSOD1 concentration (7). In this sense, although Fe^2+^ is no longer able to trigger the folding of the G93A-hSOD1, it can still bind to a set of residues of the unfolded G93A-hSOD1 ensemble (Figure 4J) to dramatically reduce the Zn^2+^-induced folding, thus blocking the first step of the G93A-SOD1 maturation (Figure 4J). As such, in the presence of Fe^2+^ at high concentration, the immature G93A-hSOD1 becomes accumulated for interacting with membranes or forming the iron-containing inclusion as observed *in vivo* (Figure 4H). The iron-induced failure of the G93A maturation will aggravate oxidative stress, which might further prompt the breakdown of blood–central nerve system barrier as a positive feedback loop. This mechanism also resolves a long-standing paradox that on the one hand, many mature ALS-causing mutants such as G93A-hSOD1 have their structures and activity highly similar to those of the mature wild-type SOD1 (13,14,28), but *in vivo*, they have been found to be highly toxic and aggregation-prone (13,14).

Our study also raises some interesting issues to be explored in the future. For example, it remains to investigate whether the Fe^2+^-bound hSOD1, which is chemically similar to Cu^+^-bound hSOD1, also acquires the activity to catalyze endogenous production of nitric oxide to induce apoptosis, as previously detected on the copper-bound, zinc-deficient SOD1 (44). Nevertheless, the high-resolution NMR results unequivocally decipher that in addition to Fenton reaction (Figure 4K), Fe^2+^ is also able to manifest its toxicity through a SOD1-dependent mechanism, by which Fe^2+^ is able to trap the mutant or even wild-type hSOD1 into highly toxic forms; as well as to provoke significant oxidative stress (Figure 4L). Both are expected to contribute to pathogenesis of neurodegenerative diseases including ALS as well as aging. This may partly rationalize the observation that under certain pathological conditions, the wild-type hSOD1 is also able to acquire cellular toxicity to initiate ALS. Furthermore, as hSOD1 exists in all human tissues, the hSOD1-dependent mechanism we decipher here is expected to play general roles in triggering other diseases upon the bio-availability of Fe^2+^ at high concentrations. Our study also provides mechanistic supports to the therapeutic approaches to treat neurodegenerative diseases or to slow down aging by repairing the breakdown of the blood-central nerve system barrier or/and intaking iron-chelators by design or from natural products such as in fruits and green tea (32-40).

## Acknowledgements

We thank Ms Linlin Miao for preparing the proteins. The study is supported by Ministry of Education of Singapore (MOE) Tier 2 Grant MOE 2011-T2-1-096 (R154-000-525-112) to Jianxing Song.

## Supplementary Materials

### Materials and methods

#### Preparation of the wild-type and mutant hSOD1proteins.

The gene encoding the wild-type hSOD1 of the native sequence was purchased from Genscript with *E. coli* preferred codons (21), while the hSOD1 mutants including G93A, H46R/H48Q and H80S/D83S were made using the QuikChange site-directed mutagenesis kit (Stratagene, La Jolla, CA). To remove the inference of extra residues, the genes of hSOD1 and their mutants were subsequently cloned into a modified vector pET28a without any tag. Then the expression vectors were transformed into and overexpressed in *Escherichia coli* BL21 (DE3) cells (Novagen). The recombinant hSOD1 and mutant proteins were found in inclusion body. As a result, the pellets were first washed with buffers several times and then dissolved in a phosphate buffer (pH 8.5) containing 8 M urea and 100 mM dithiothreitol (DTT) to ensure complete conversion to Cys-SH. After 1 hour, the solution were acidified by adding 10% acetic acid and subsequently purified by reverse-phase (RP) HPLC on a C4 column eluted by water-acetonitrile solvent system. The HPLC elutions containing pure recombinant hSOD1 and mutants were lyophilized and stored at -20 degree.

The generation of the isotope-labeled proteins for NMR studies followed a similar procedure except that the bacteria were grown in M9 medium with the addition of (^15^NH_4_)_2_SO_4_ for ^15^N labeling and (^15^NH_4_)_2_SO_4_ /[^13^C]-glucose for ^15^N-/^13^C-double labelling (21). The purity of the recombinant proteins was checked by SDS–PAGE gels and verified by a Voyager STR matrix-assisted laser desorption ionization time-of-flight-mass spectrometer (Applied Biosystems), as well as NMR assignments. The concentration of protein samples was determined by the UV spectroscopic method in the presence of 8 M urea (21).

#### NMR experiments.

As the hSOD1 of the native sequence is high prone to aggregation, we thus have screened a variety of solution conditions and successfully found the optimized conditions as facilitated by our previous studies on insoluble proteins (21,25,45). All wild-type and mutants hSOD1 samples were prepared in milli-Q water at pH 3.5 because: 1) this condition allows the preparation of soluble and stable hSOD1 samples with a concentration up to 500 μM, which is required to collect high-quality triple-resonance NMR spectra for assigning the folded hSOD1 weakly-populated in the presence of Zn^2+^. 2) As we previously found (21), the hSOD1 of the native sequence containing four cysteine residues, started to form both native and nonnative disulfide bridges even in the presence of DTT once the solution pH is above 4.0, and consequently hSOD1 becomes partially folded (21). This unusual property of hSOD1 of the native sequence is reflected by its ability to form and maintain the disulfide bridge in the reducing cytosolic environment. 3) At this pH, the solubility of Fe^2+^ is high enough to allow our titrations to reach high ratios of Fe^2+^.

All NMR experiments were acquired at 25 degree on an 800 MHz Bruker Avance spectrometer equipped with pulse field gradient units as described previously (21,24,25). For characterizing the unfolded hSOD1 ensemble, a pair of triple-resonance experiments HNCACB, CBCA(CO)NH were collected for the sequential assignment on a ^15^N-/^13^C-double labelled sample of 500 μM, while ^15^N-edited HSQC-TOCSY and HSQC-NOESY were collected on a ^15^N-labelled sample at a protein concentration of 500 μM. For achieving assignments of the folded hSOD1 induced by Zn^2+^, triple-resonance experiments HNCACB, CBCA(CO)NH were collected for on a ^15^N-/^13^C-double labelled sample of 500 μM in the presence of Zn^2+^.

For assessing the backbone dynamics of the unfolded hSOD1 ensemble on the ps-ns time scale, {^1^H}-^15^N steady-state NOEs were obtained by recording spectra on the ^15^N-labeled sample at 500 μM with and without ^1^H presaturation with duration of 3 s plus a relaxation delay of 6 s at 800 MHz. To assess conformational exchanges over μs-ms, ^15^N transverse relaxation dispersion experiments were acquired on the ^15^N-labeled sample on a Bruker Avance 800 spectrometer with a constant time delay (TCP = 50 ms) and a series of CPMG frequencies, ranging from 40 Hz, 80 Hz, 120 Hz (x3), 160 Hz, 200 Hz, 240 Hz, 320 Hz, 400 Hz, 480 Hz, 560 Hz, 640 Hz, 720 Hz, 800 Hz, and 960 Hz (×3 indicates repetition) as we previously performed (21,24).

## Supplementary Figures

**Fig. S1.**
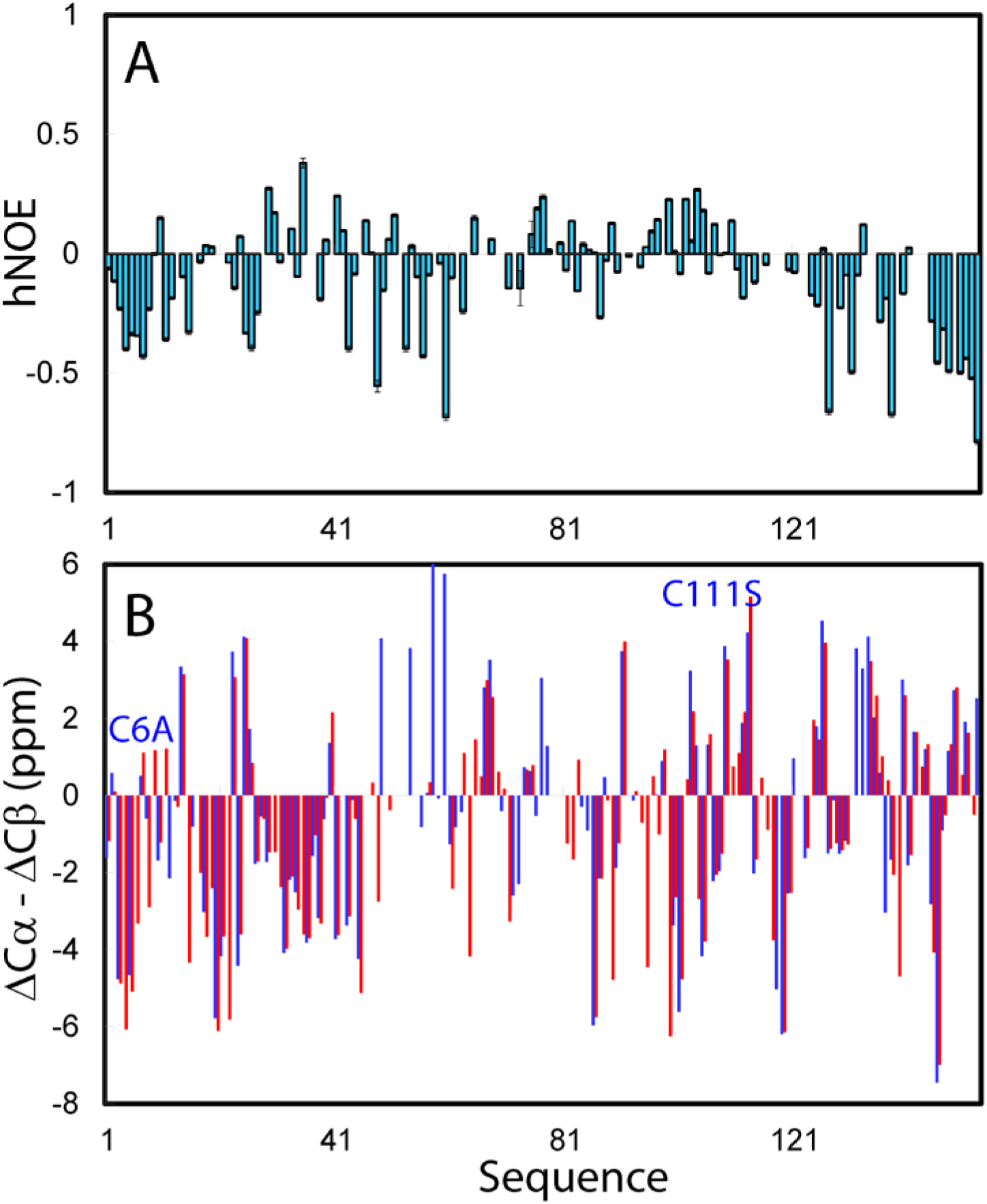
NMR characterization of residue-specific conformations and dynamics of the unfolded and folded hSOD1. (A) {^1^H}-^15^N heteronuclear steady-state NOE (hNOE) of the unfolded hSOD1 without metallation and disulfide bridge, which mimic nascent hSOD1. (B) Residue specific (ΔCα-ΔCβ) chemical shifts of the Zn^2+^-bound hSOD1 of the native sequence (red), and those derived from the previously deposited NMR data (BMRB Entry 6821) for the Zn^2+^-bound hSOD1 with C6A/C111S mutations (blue). The two mutation sites C6 and C111 are indicated.

**Fig. 2.**
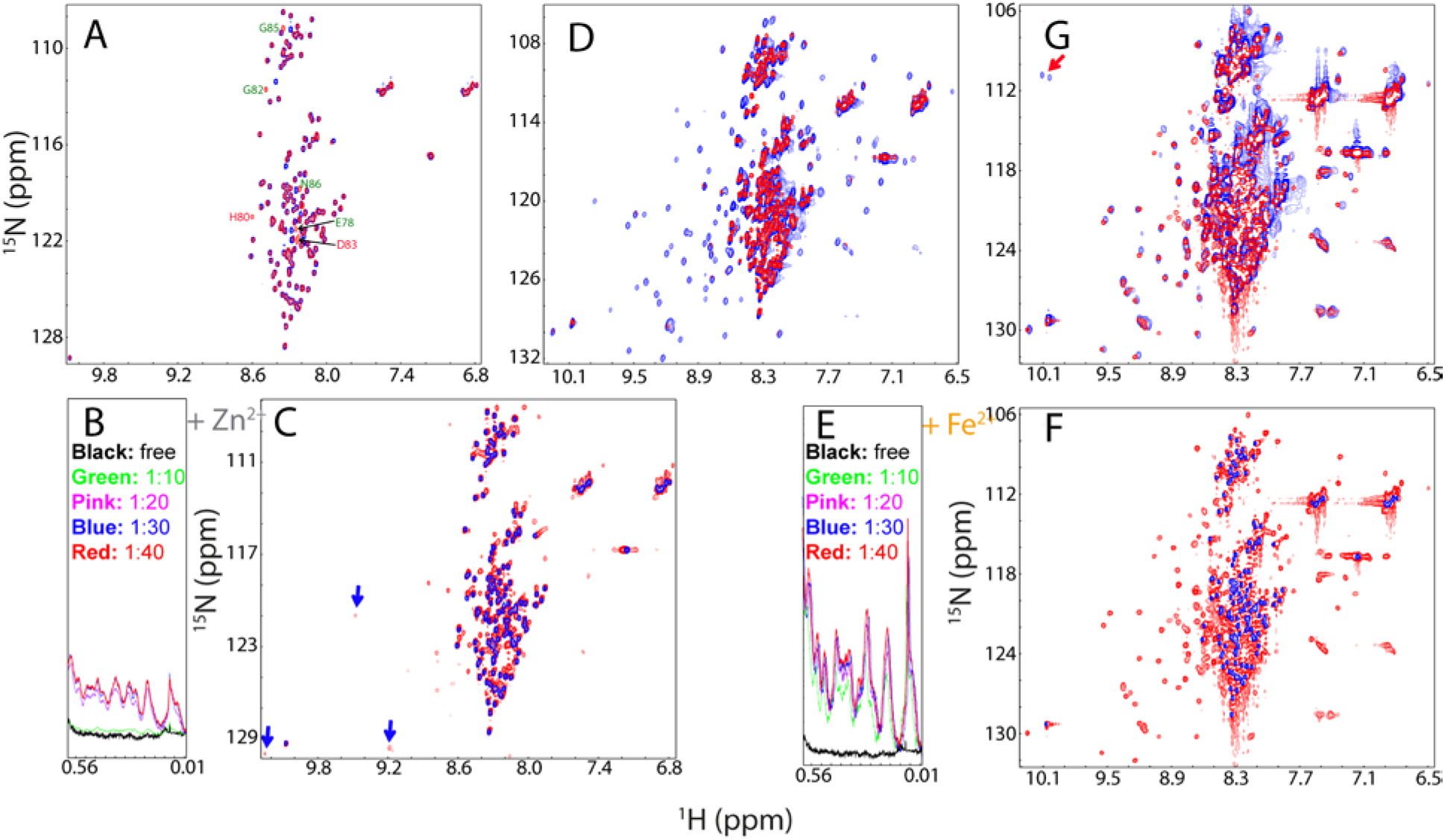
NMR characterization of interactions of the H80S/D83S hSOD1 with Zn^2+^ and Fe^2+^. (A) Superimposition of HSQC spectra of the wild-type hSOD1 (red) and H80S/D83S hSOD1 (blue) without metalation and disulfide bridge. Residues with significant difference for their HSQC peaks are labeled. (B) Up-field 1D NMR peaks characteristic of the folded form of H80S/D83S hSOD1 (0.0-0.6 ppm) induced by the presence of Zn^2+^ at different molar ratios. (C) Superimposition of HSQC spectra of H80S/D83S hSOD1 in the absence (blue) and in the presence of Zn^2+^ at a molar ratio of 1:40 (red). Some well-dispersed peaks characteristic of the partially-folded form are indicated by green arrows. (D) Superimposition of HSQC spectra of the wild-type hSOD1 in the presence of Zn^2+^ at a molar ratio of 1:20 (blue), and H80S/D83S hSOD1 in the presence of Zn^2+^ at a molar ratio of 1:40 (red). (E) Up-field 1D NMR peaks characteristic of the folded form of H80S/D83S hSOD1 (0.0-0.6 ppm) in the presence of Fe^2+^ at different molar ratios. (F) Superimposition of HSQC spectra of H80S/D83S hSOD1 in the absence (blue) and in the presence of Fe^2+^ at a molar ratio of 1:20 (red). (G) Superimposition of HSQC spectra of the wild-type hSOD1 (blue) and H80S/D83S hSOD1 (red) in the presence of Fe^2+^ at a molar ratio of 1:20.

**Fig. S3.**
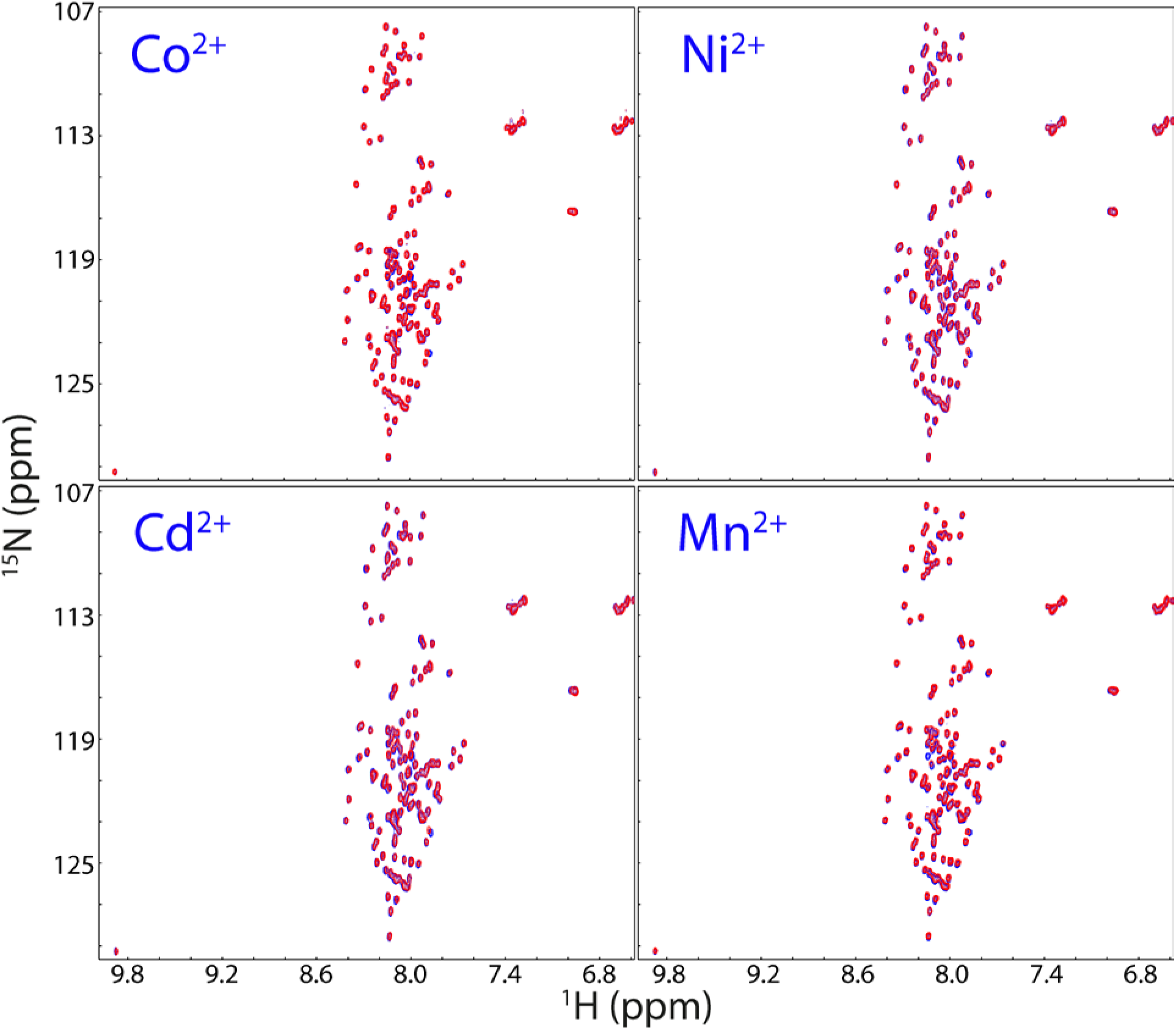
HSQC characterization of interactions of the unfolded hSOD1 with four cations. Superimposition of HSQC spectra of hSOD1 without metalation and disulfide bridge (blue) and in the presence of Co^2+^, Ni^2+^, Cd^2+^, and Mn^2+^ respectively at a molar ratio of 1:40 (red).

**Fig. S4.**
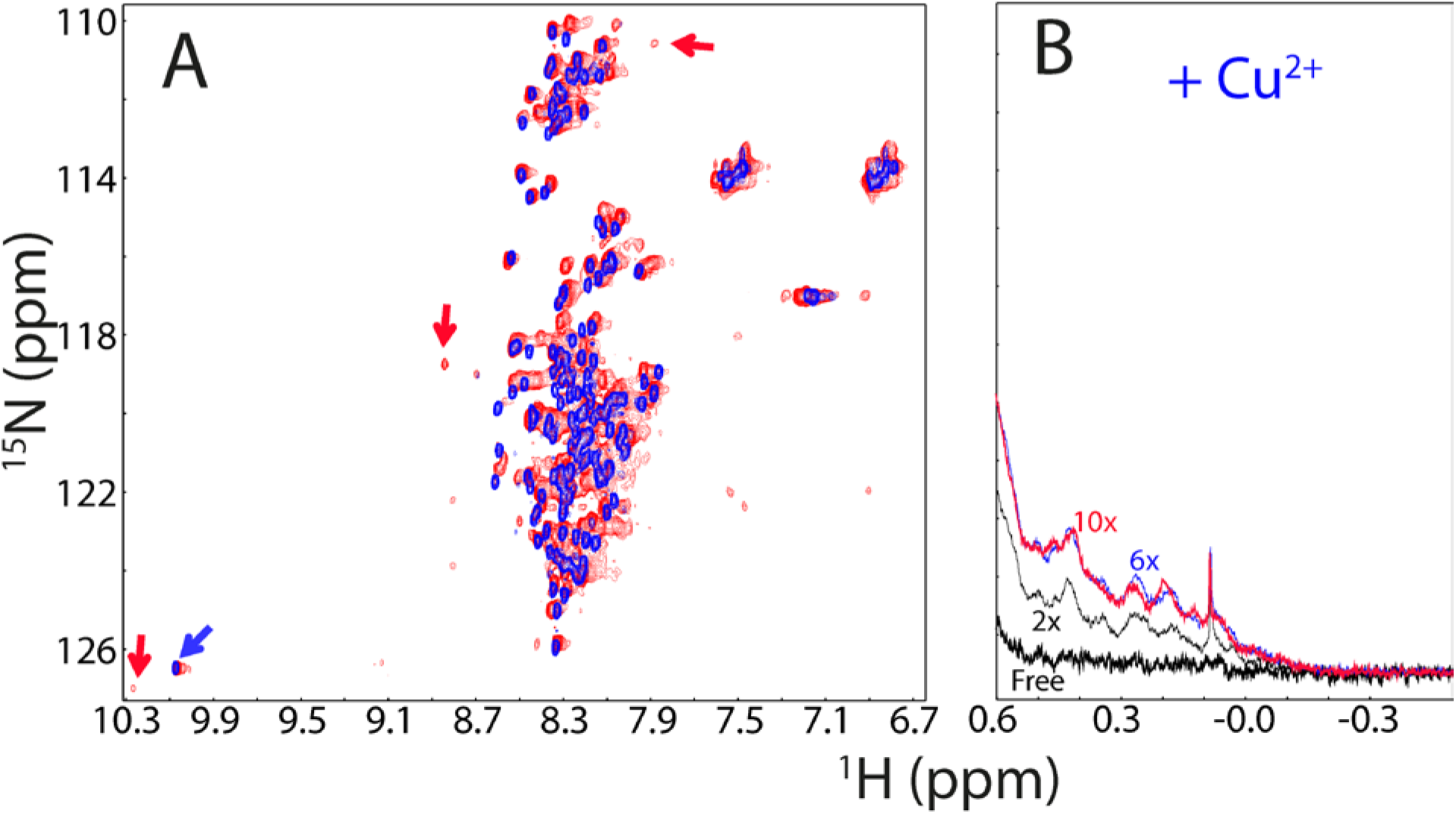
Copper has extensive binding to the unfolded hSOD1 ensemble. (A) Superimposition of HSQC spectra of hSOD1 without metalation and disulfide bridge in the absence (blue) and in the presence of Cu^2+^ at a molar ratio of 1:10 (red). The green arrow is used for indicating the HSQC peak of Trp32 ring proton characteristic of the unfolded ensemble and red ones for some HSQC peaks characteristic of the partially-folded form. (B) The up-field 1D NMR peaks characteristic of the partially-folded hSOD1 (-0.5-0.62 ppm) in the presence of Cu^2+^ at different molar ratios.

**Fig. 5.**
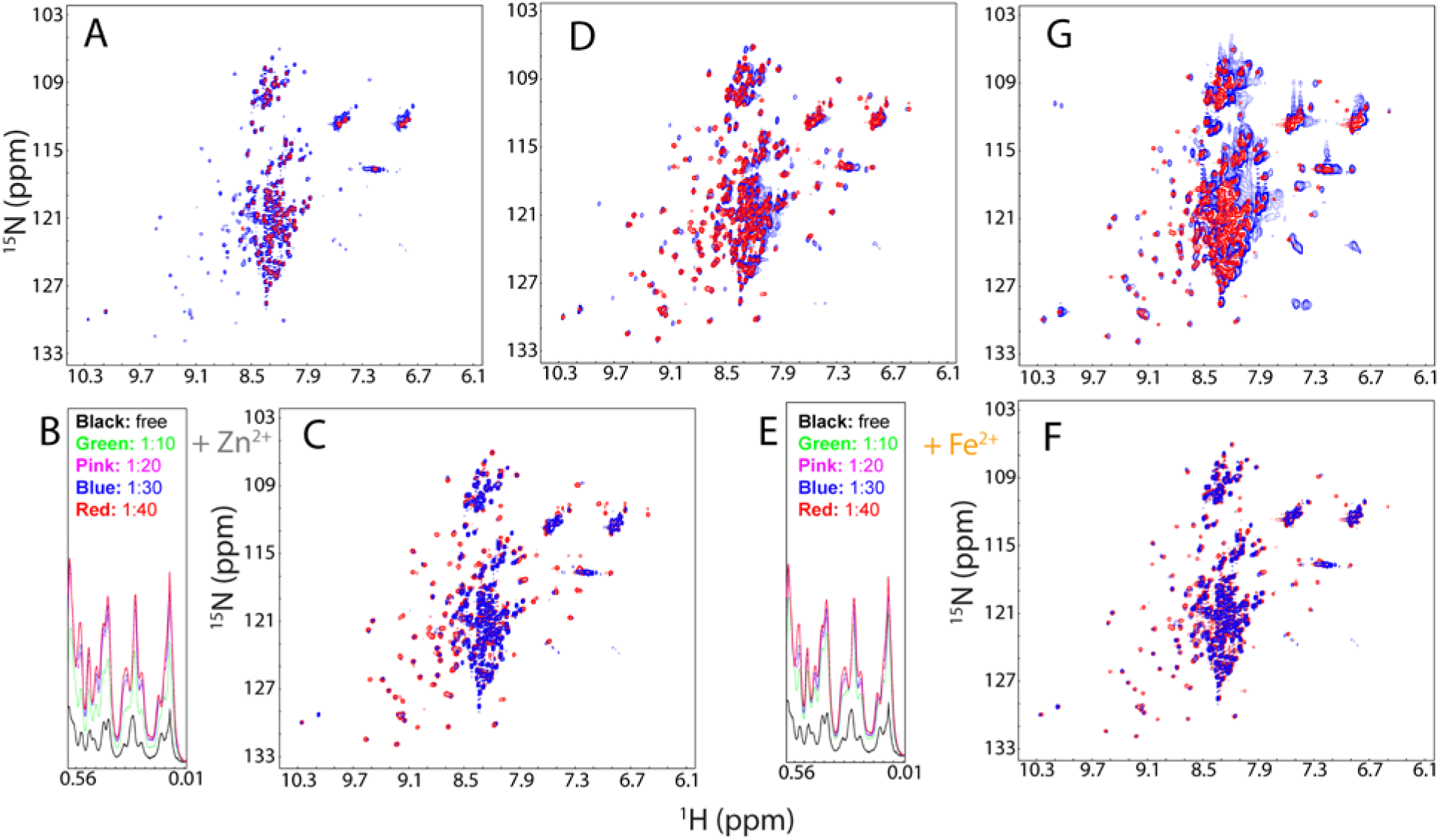
NMR characterization of interactions of the H46R/H48Q hSOD1 with Zn^2+^ and Fe^2+^. (A) Superimposition of HSQC spectra of the wild-type hSOD1 (red) and H46R/H48Q hSOD1 (blue) without metalation and disulfide bridge. (B) Up-field 1D NMR peaks characteristic of the folded form of H46R/H48Q hSOD1 (0.0-0.6 ppm) in the presence of Zn^2+^ at different molar ratios. (C) Superimposition of HSQC spectra of H46R/H48Q hSOD1 in the absence (blue) and in the presence of Zn^2+^ at a molar ratio of 1:40 (red). (D) Superimposition of HSQC spectra of the wild-type hSOD1 in the presence of Zn^2+^ at a molar ratio of 1:20 (blue), and H46R/H48Q hSOD1 in the presence of Zn^2+^ at a molar ratio of 1:40 (red). (E) Up-field 1D NMR peaks characteristic of the folded form of H46R/H48Q hSOD1 (0.0-0.6 ppm) in the presence of Fe^2+^ at different molar ratios. (F) Superimposition of HSQC spectra of H46R/H48Q hSOD1 in the absence (blue) and in the presence of Fe^2+^ at a molar ratio of 1:20 (red). (G) Superimposition of HSQC spectra of the wild-type hSOD1 (blue) and H46R/H48Q hSOD1 (red) in the presence of Fe^2+^ at a molar ratio of 1:20.

**Fig. S6.**
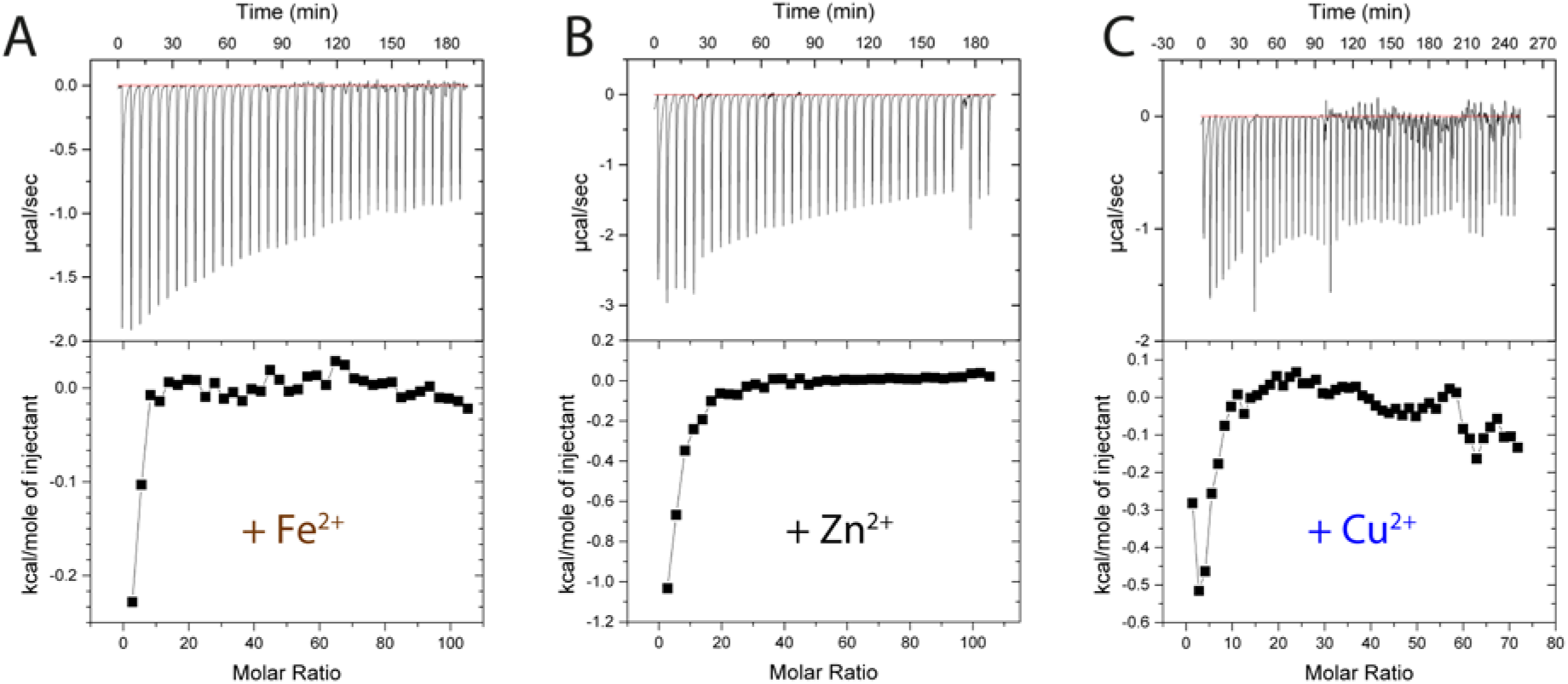
ITC characterization of interactions of the unfolded hSOD1 with three cations. (Upper) ITC profiles of the interactions of the unfolded hSOD1 ensemble with Fe^2+^ (A); Zn^2+^ (B) and Cu^2+^ (C), and (Lower) integrated values for reaction heats with subtraction of the corresponding blank results normalized against the amount of ligand injected versus the molar ratio of hSOD1:cation.

**Fig. S7.**
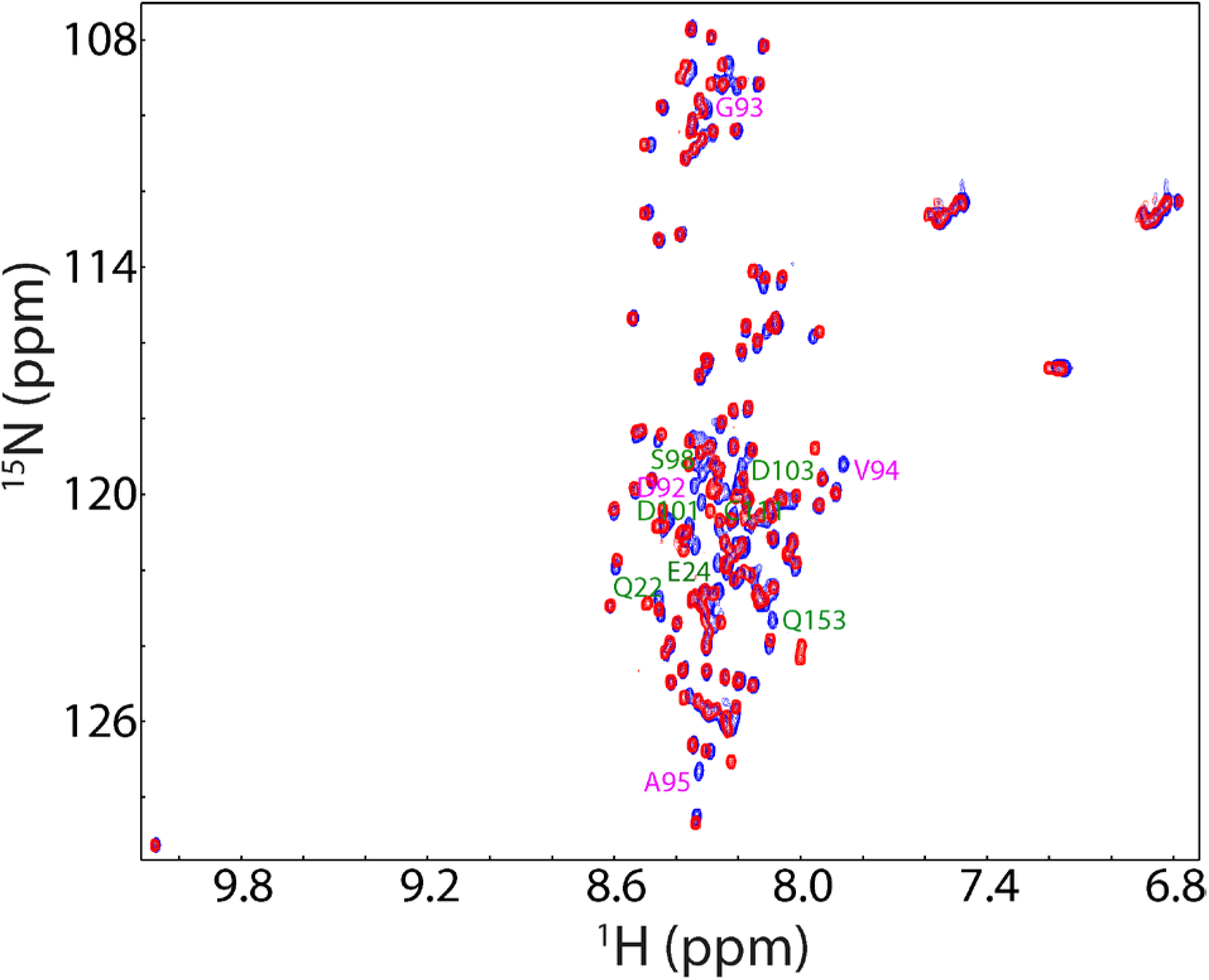
HSQC characterization of G93A hSOD1. Superimposition of HSQC spectra of the wild-type (blue) and G96A hSOD1 (red) without metalation and disulfide bridge. Residues with significant shifts for their HSQC peaks are labeled.

**Fig. S8.**
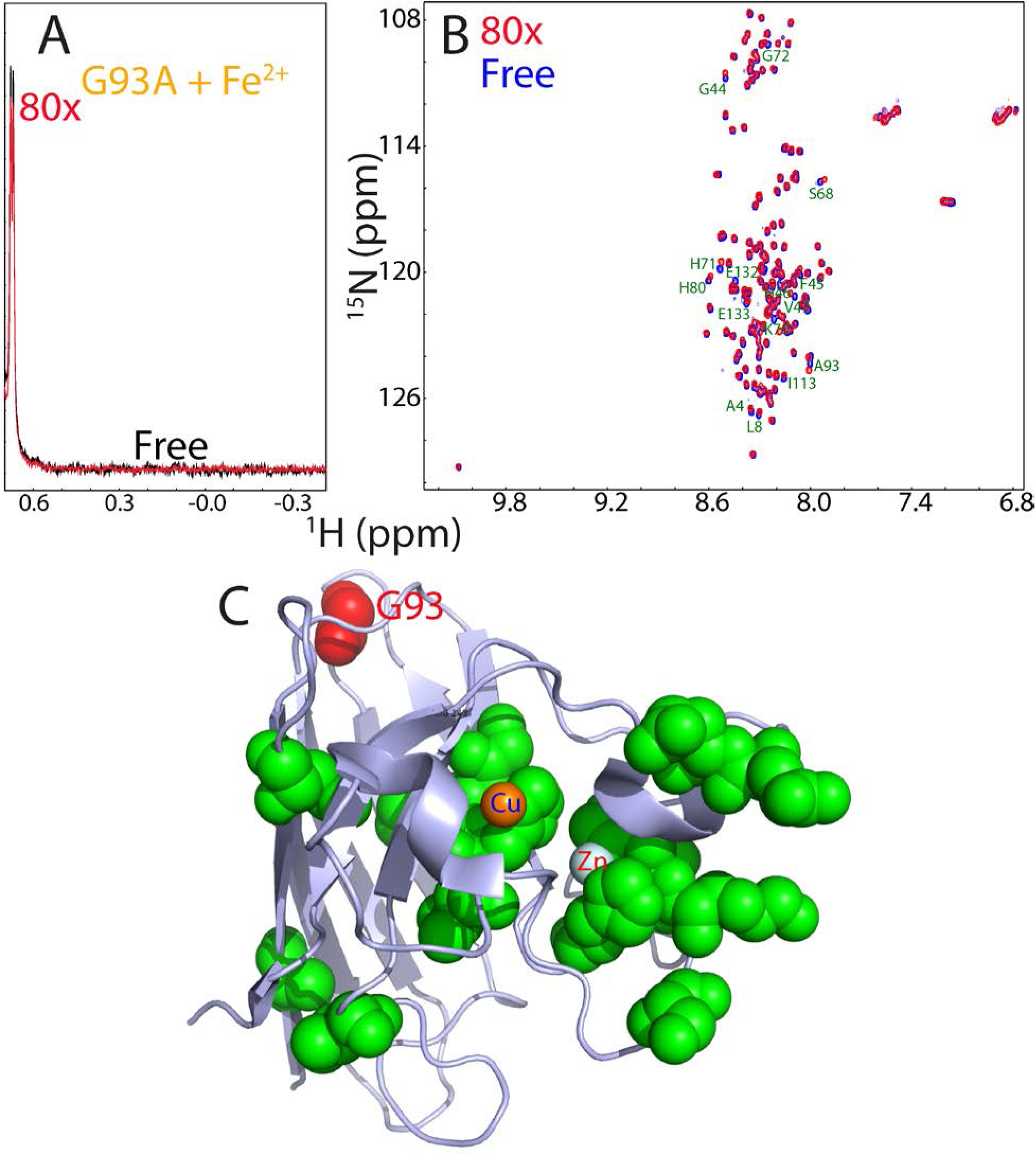
G93A hSOD1 loses the capacity of folding as induced by Fe^2+^. (A) Up-field NMR peaks (-0.5-0.64 ppm) of G93A hSOD in the absence (black) and in the presence of Fe^2+^ at a molar ratio of 1:80 (red). (B) Superimposition of HSQC spectra of G93A hSOD1 in the absence (blue) and in the presence of Fe^2+^ at a molar ratio of 1:80 (red). Residues with significant shifts for their HSQC peaks are labeled.

**Fig. S9.**
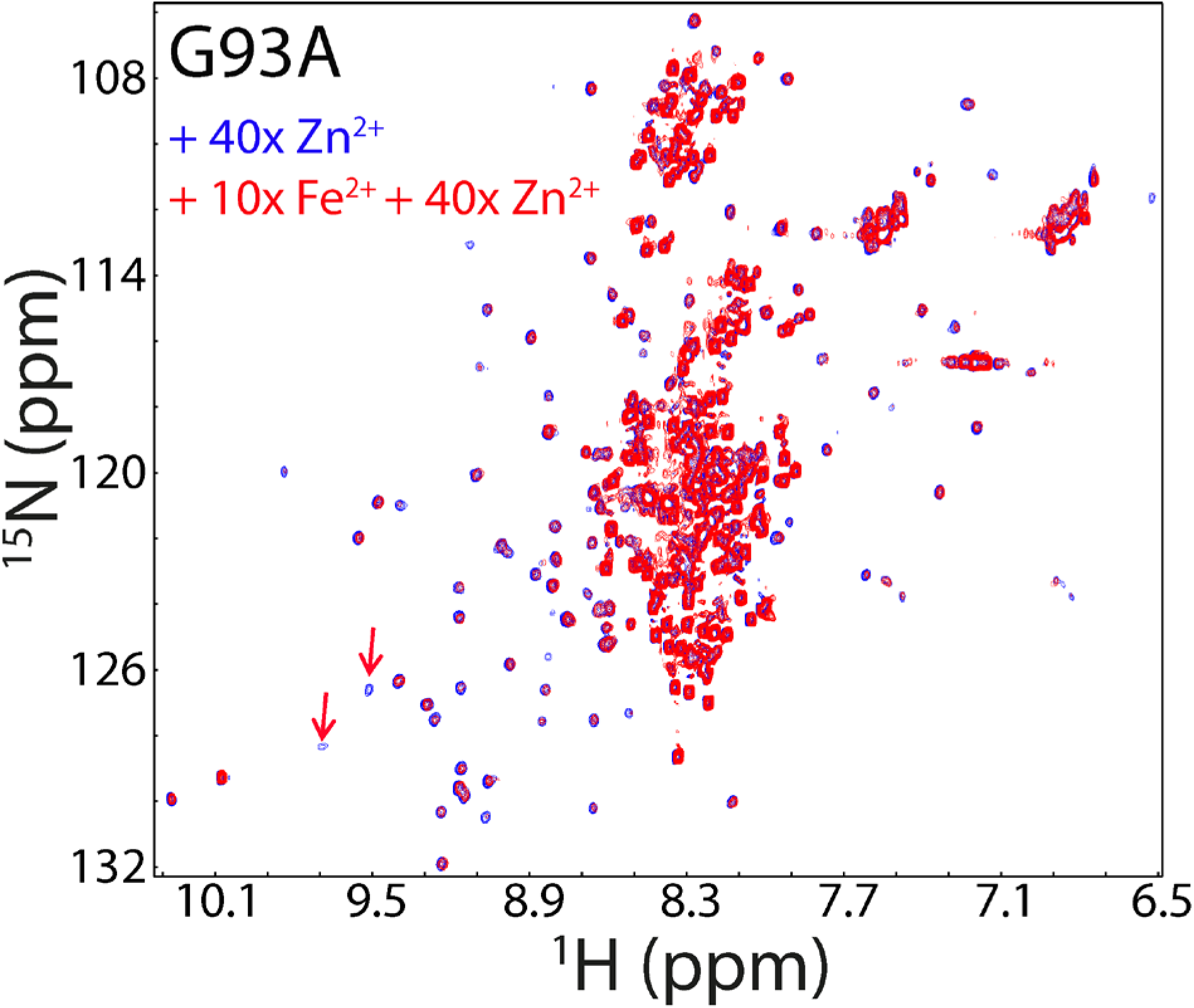
The pre-existence of Fe^2+^ significantly reduces the zinc-induced folding of G93A hSOD1. Superimposition of HSQC spectra of G93A hSOD1 in the presence of Zn^2+^ at a molar ratio of 1:40, without (blue) and with the pre-existence of 10x Fe^2+^ (red). Red arrows are used to indicate some well-dispersed HSQC peaks characteristic of the folded form which are too weak to be detected with the pre-existence of 10x Fe^2+^.

